# Modular Arrays for High Precision Wearable MEG

**DOI:** 10.64898/2026.08.19.745485

**Authors:** Nicholas A. Alexander, Alberto Mariola, Sahitya Puvvada, Yulia Bezsudnova, Tim M. Tierney, Gareth R. Barnes, Martina F. Callaghan

**Affiliations:** Functional Imaging Laboratory (FIL), Department of Imaging Neuroscience, UCL Queen Square Institute of Neurology, University College London, London, UK

**Author notes:** **Corresponding author:** Nicholas A. Alexander.

**Keywords:** OPM, wearable, MEG, mobile, naturalistic, ambulatory

## Abstract

Optically pumped magnetometers (OPMs) can be used for magnetoencephalography (MEG) with equivalent or improved signal to noise ratio, relative to cryogenic MEG, when sensors are placed close to the scalp. OPM-based MEG can also be used in mobile contexts if sensors are placed in lightweight, wearable arrays. Individually tailored, rigid helmets known as scannercasts are currently the only method capable of achieving on-scalp, mobile recordings with high precision. However, these scannercasts are expensive to produce, require structural imaging in advance of the experiment, and can incur lengthy downtime while sensors are transferred between scannercasts. Here, we introduce a solution to these challenges that retains the advantages of scannercasts. We provide detailed steps for constructing a modular, cap-based design, suitable for all head sizes. Using simulations, we compare the leadfield power of this array against an idealised array and a commercially available mobile solution. We then validate our proposed solution empirically, in five participants, and provide a complete data preparation and analysis pipeline. Our design expands the accessibility of OPM-based MEG, and increases participant throughput to levels comparable to other imaging modalities. Crucially, it removes the trade-off between signal quality, mobility and practicality, promoting the unique potential of OPM-based MEG as a tool for studying naturalistic behaviour, and clinical assessment with high precision.

## 1. INTRODUCTION

Optically pumped magnetometers (OPMs) have given researchers freedom to design their own systems for optically pumped magnetoencephalography (MEG; OP-MEG) due to their small size and capacity for flexible positioning. Early work used small arrays positioned over select regions of interest (Barry et al., 2019; Boto et al., 2017; Lin et al., 2019). However, the commercial availability of OPM hardware engineered specifically for MEG, has made whole-head systems containing 64-128 sensors increasingly common (Alem et al., 2023; Bonnet et al., 2025; Schofield et al., 2024). A fundamental challenge remaining for the field is to devise a means of positioning the OPM sensors as close to the scalp as possible, to maximise signal amplitude and the precision of source localisation, but with a generic configuration that is not bespoke to the individual.

Existing designs can be broadly categorised as static versus mobile, and one-size-fits-all versus truly on-scalp. Static systems mimic the constraints of superconducting quantum interference device (SQUID) based MEG systems. Participants cannot move their head and the distance between sensors and scalp increases at the extremities of the array, with a corresponding reduction in signal-to-noise ratio (SNR) (Boto et al., 2016; Iivanainen et al., 2017). In contrast, mobile, participant-specific designs place sensors directly on the scalp, ensuring consistently minimal sensor-scalp separation and allowing free movement (Mellor et al., 2023; Seymour et al., 2021).

Static solutions exist, which allow radial translation of sensors to fit a wide range of head shapes (Alem et al., 2023; Hristova et al., 2026; Schwartz et al., 2025), offering a cryogen-free alternative to SQUID-MEG. On the other hand, mobile systems enable a broader range of experimental paradigms that would be infeasible with static MEG, e.g. measuring neural activity during naturalistic behaviours such as dance (O’Neill, Seymour, et al., 2025), locomotion (Seymour et al., 2021; Spedden et al., 2025) and social interaction (Holmes et al., 2023).

Scannercasts are rigid, participant-specific helmets constructed from scalp models typically produced using structural magnetic resonance imaging (MRI) (Boto et al., 2017; Troebinger et al., 2014). In addition to the benefit of sensor-to-scalp proximity, scannercasts provide sensor stability on the head during movement (Seymour et al., 2021), automatic spatial co-registration to anatomy, and consistency of sensor positioning across sessions (Meyer et al., 2017). However, scannercasts are difficult to scale for OP-MEG due to the costs associated with multiple visits and manufacturing, and the downtime while sensors are transferred between scannercasts limiting participant throughput. Assumed automatic spatial co-registration is also sensitive to deformation and manufacture error (Hill et al., 2025; Pang et al., 2022).

Another approach is to use generic helmets designed to accommodate a range of head shapes and select the size that best fits the individual (Gao et al., 2023; Schofield et al., 2024). Having multiple generic helmet sizes is effective at reducing the effect of head size on scalp-to-sensor distance compared with a true one-size-fits-all approach (Rhodes et al., 2023; Rier et al., 2024). However, like scannercasts, this requires moving sensors from helmet to helmet. In addition, the benefits of automatic co-registration are lost and sensor placement is not optimised for the individual. Nonetheless, generic helmets are effective at facilitating large cohort studies (Rivero et al., 2025; Safar et al., 2025), and can be used during movement (Sanders et al., 2025; Schofield et al., 2024).

Flexible, cap-based systems common in mobile electroencephalography (EEG) and functional near infra-red spectroscopy (fNIRS) offer an alternative approach. Caps are available in multiple sizes and use mechanical fixtures to secure equipment (e.g. electrodes, optodes). Similar cap-based approaches have been used for OPMs (Anders et al., 2026; Cao et al., 2023; Feys et al., 2025; Hill et al., 2020; Power et al., 2024), but with known limitations related to sensor movement and transfer time, which we aimed to address.

In this article, we present a cap-based solution for mounting modules containing multiple sensors, enabling on-scalp, mobile MEG at low cost, with high throughput and minimal downtime. We first describe the design plan and implementation, with steps to reproduce the system provided. Next, in simulation we verify the signal quality of this approach compared with an idealised array and an existing generic helmet design. Finally, we empirically demonstrate successful application of our module array in five participants. We anticipate this modular, cap-based solution will improve accessibility and throughput of OP-MEG, as required for widespread clinical and cognitive neuroscience applications, particularly where both freedom of movement and high precision are required.

## 2. MATERIALS AND METHODS

To enable multiple scanning sessions per day, the time required to transfer sensors between caps must be minimal, as must the time taken to apply the sensors to the participant. To guard against population biases, the system must not exclude participants based on age, weight, head shape or hairstyle. We therefore aimed to produce a setup that would easily enable sensors to be inter-changed between caps, which themselves could be placed on the head quickly, and would conform snugly and comfortably to the surface of the head.

It is also important to be compatible with the range of experiments conducted using OPMs. These include imaging of the spinal cord (Mardell et al., 2024) and heart (Li et al., 2025; Woelk et al., 2026) and combination with head-mounted displays (Roberts et al., 2019; Zabbah et al., 2026). A secondary aim was therefore to produce a solution which can, in future, be adapted to, and integrate with, a variety of imaging configurations beyond the brain.

### 2.1. OP-MEG System

Our modular, cap-based system has been designed for a QuSpin Neuro-1 system (QuSpin Inc., CO, USA) comprising 64 tri-axial sensors (192 channels) (Schofield et al., 2024). The Neuro-1 houses sensor control electronics inside a wearable backpack, and bundles of eight sensors connect to circuit boards via lightweight, 1.5 mm diameter round cables of ∼2m length.

Although developed around the QuSpin OPM architecture, the design principles described here are applicable to other OPMs with similar housing dimensions (approximately 42 x 17 x 13 mm), such as Fieldline sensors (Fieldline Inc. CO, USA), enabling simple adaptation to alternative systems.

### 2.2. Design and Manufacturing

#### 2.2.1 Design Objectives

A mobile system for OP-MEG which is scalable and accessible must meet several requirements. Primarily, it must: (i) achieve signal amplitude measurement that is at a level comparable to existing methods; (ii) maintain consistent sensor-source positioning during participant movement; and (iii) support high participant throughput.

Signal amplitude of sources in the brain is dependent on the density, proximity and coverage of the array, in addition to the noise conditions of the sensor and the field measurement configuration (Boto et al., 2016; Schoffelen et al., 2025). We aimed to design a system with whole head coverage and proximate, equidistant spatial sampling.

During movement it is essential that sensors do not move with respect to each other, or the head. This is to maintain spatial co-registration and to minimise vibration of sensors which would introduce sensor-dependent interference that is difficult to model (Ferez et al., 2026). Further, random co-registration error has been demonstrated to have greater negative effect on source reconstruction when the error is sensor-specific rather than being consistent for the whole array (Hill et al., 2020), although this effect is reduced by scalp proximity (Iivanainen, 2026). We aimed to mount sensors in a low-profile, large footprint arrangement that minimises the forces enacted on individual sensors during movements and keeps them close to the scalp.

#### 2.2.2. Module Design

All computer-aided design (CAD) was completed using the open-source software Blender (Blender Foundation, Amsterdam, Netherlands).

We designed rigid modules that each contain four sensors (12 channels) positioned on their side (**Figure 1A**). Grouping sensors into shared modules supports several of our design goals. First, it reduces the number of installation steps from one per sensor to one per module (e.g. 16 operations instead of 64), which reduces transfer time between caps. The number of independent cables to manage is also reduced, saving time. Similar design principles were used in the Kernel Flux OP-MEG system which arranged nine sensors within a single housing (Pratt et al., 2021). Second, the module housing protects the sensors from direct handling increasing their robustness, and providing heat insulation. Third, the larger footprint of a module increases stability during movement. Finally, orienting sensors on their sides lowers the radial profile of the array, reducing forces on the sensors during head motion.

**Fig. 1.**
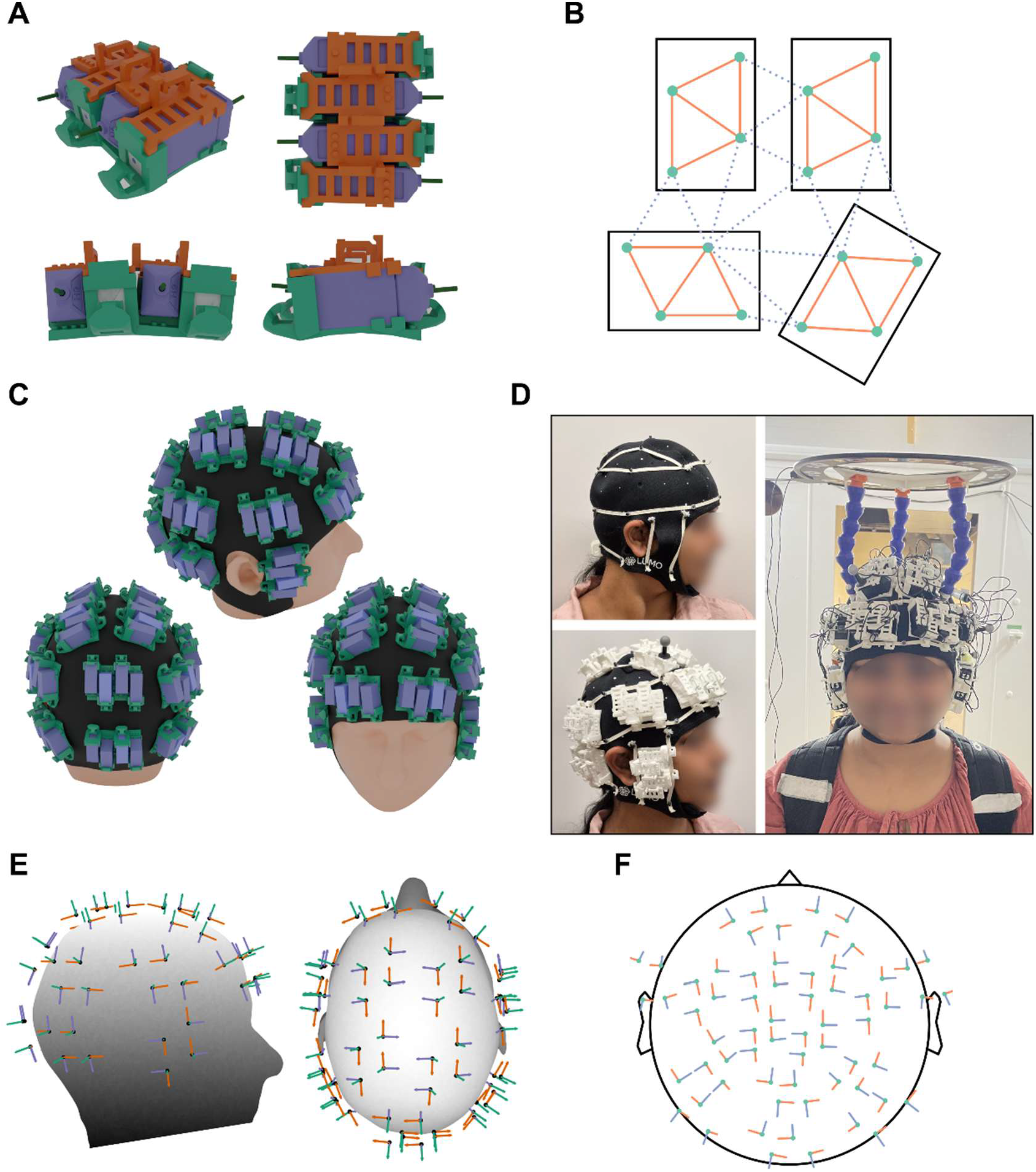
Sensor module design. (A) Digital render of the sensor modules from multiple angles. The base is shown in green, the lid with cable supports in orange and sensors in purple. The 10° curvature can be seen in the lower panels. Note, colours shown do not match real production. (B) Drawing of sensor module footprint (solid black lines) with the sensitive parts of the cell in each sensor in a module marked by green circles. Approximately equilateral triangles with ∼30 mm edges are formed between green circles within modules (solid orange lines) and, under ideal conditions (top), between nearby modules (dashed blue lines). Also shown (bottom) are realistic placements of modules where between-module distances are greater than 30 mm. (C) Digital render of dense, non-overlapping placement of sensor modules onto an adult male head (∼56 cm circumference). In this arrangement, 64 sensors (purple) can be attached. Note, the module lid and OPM cables are not shown, to better illustrate module and sensor footprint. (D) Photographs of a co-author of this article (with permission) wearing the system within the MSR. The cotton elastic fixtures can be seen in the top left and the module placement in the bottom left. The complete setup, including the flexible HALO mounting system is shown on the right. (E) Field measurement positions and orientations for the 64-sensor layout shown in C. (F) Projection of sensor positions and orientations shown in C-D into 2D.

Using modules does introduce several challenges. The head is variably curved and so a large, flat module would sit unevenly and increase sensor–scalp distance. Modules must also avoid overlapping while still providing full coverage across a large range of head sizes. One way to mitigate these issues is to minimise module size while maximising internal distances between magnetometer locations. This also allows external distances between neighbouring modules to be reduced, supporting equidistant coverage (**Figure 1B-C**). High SNR can be achieved with approximately equidistant spatial sampling of ∼30 mm (Ahonen et al., 1993; Tierney et al., 2020), so we arranged the sensors within modules to form two approximately equilateral triangles, each with ∼30 mm edges. We oriented the sensors so that the sensitive parts of the cells lay at the triangle vertices. Positioning the sensitive region toward the external edge of the module further reduced its overall size.

Another design constraint is the size and orientation of the sensor housing and the offset between the housing surface and the sensitive location of the cell (i.e., the field-measurement point). Orientating QuSpin Neuro-1 QZFM sensors on their side, with the X axis radial to the scalp, results in a minimum 6.25 mm distance to the scalp. We made the base of the module 2.1 mm thick beneath the sensor. Combined with the approximately 3 mm cap thickness, this results in an ∼11 mm sensor-scalp offset. We also applied incremental 5° rotations to the sensor slots, yielding ∼10° curvature across the module’s contact plane to better adhere to head curvature, noting that the neoprene cap absorbs some of the difference between head and module curvature.

Stability during movement ultimately depends on how the modules attach to the cap. The mounting system must be comfortable, adaptable to different head shapes, secure, and quick to use. These requirements preclude rigid fixtures common when mounting single sensors (Feys et al., 2025; Hill et al., 2020). We therefore used 5 mm elastic cotton tape attached under tension to form bands across the cap (**Figure 1D**). Each module has two hooks on its long edges that engage with these bands. The interlocking design allows tension to be shared between adjacent modules and enables close spacing. Once compressed by the elastic, the module beds into the neoprene surface to form a stable mount. Modules can be attached before the cap is placed onto the head which reduces setup time with participants.

#### 2.2.3. Cap Design

Across fNIRS and EEG, a variety of commercial cap solutions exist. We selected LUMO (Gower Labs Ltd., London, UK) neoprene caps, originally developed for fNIRS, based on our requirements for whole-head coverage (i.e. extending from nasion to inion), heat insulation and stability. These caps are approximately 3 mm thick and available for head sizes from 34 to 64 cm in circumference, in 2 cm increments, allowing for a close fit across child and adult head sizes. Other designs (e.g. Artinis Headcap, Artinis Medical Systems, NL) or bespoke caps could also be viable solutions, though only the LUMO cap has been tested here. Our choice of an fNIRS cap reflects similarities in requirements, in particular regarding the weight and stability of sensors (Vidal-Rosas et al., 2023).

Neoprene is well suited to this application due to its elasticity, elastic recovery, and tensile strength. It has widespread use in applications such as wetsuits where flexibility and durability are critical, as well as offering biocompatibility for skin contact. Neoprene also permits robust attachment of hardware. Modules can be secured either by sewing with nylon thread or by punching holes to accept inserts, such as nylon cinch rivets. We opted to use the latter as it allows precise placement and parts can be replaced easily.

This approach also meets our secondary design aim because it can be readily adapted to other regions of interest that can be imaged by measuring magnetic fields, e.g. the spinal cord and heart (Li et al., 2025; Mardell et al., 2024). Conforming neoprene clothing is available and would allow dense arrays of OPMs to be mounted on the chest, back and neck, providing adequate coverage (O’Neill, Spedden, et al., 2025). A prototype of our module has already been used in combination with a scannercast to image heart function for the purposes of modelling and removing cardiac field artefact from MEG (Woelk et al., 2026), and future work will continue in this area.

#### 2.2.4. Cable Management

The QuSpin Neuro-1 system backpack connects via two lightweight cables to a power supply and acquisition hardware housed outside the MSR. These allow large movements like stepping (Spedden et al., 2025). However, every OPM sensor connects to the backpack via its own cable. In our case this results in 64 lightweight cables which must be protected and stabilised during movement. To achieve this, on top of each module are three loops which receive cables in a woven fashion. They can secure the four cables of sensors within the module, as well as from other modules. To provide a further anchor, the complete bundle of cables exiting the cap is secured to a clip attached at the rear of the cap, which receives cables bound with a hook-and-loop tie. This plastic clip fastens either side of the neoprene fabric and is bolted through for security. This allows for cable management to be completed before the participant arrives such that the cap can simply be placed on the head with all sensors already attached via the modules.

#### 2.2.5. Sensor Module Manufacturing

Additive manufacturing (3D printing) provides an efficient way to produce small batches of geometrically complex components without the costs associated with moulds or tooling, making it well suited to our sensor modules. We underwent several rounds of prototyping, including testing different additive manufacturing methods, materials and designs. The modules incorporate curved and recessed features and must be both strong and precise.

Ultimately, we used selective laser sintering (SLS) of PA 2200 (white nylon PA12 powder) material as it meets these requirements and has previously been used to manufacture OPM scannercasts, demonstrating compatibility with the OPM sensors. SLS printers are large and expensive, and typically require printing multiple components in a single build, and therefore likely require either institutional support or collaboration with commercial manufacturers.

A key consideration was manufacturing tolerances of both sensor housing and our modules. By using thin, flexible walls and textured surfaces we achieved a tool-free, friction fit. Note, these small variations can be addressed during calibration of array geometry.

### 2.3. Simulations

To assess the performance of our setup at capturing signals across the brain, we evaluated the leadfield power of different array configurations using simulation methods. In particular, we wanted to assess how inconsistent inter-sensor distances and sensor-scalp offsets each impacted leadfield power, as these were fundamental to our module design.

Therefore, simulations were performed using three sensor array configurations: our proposed modular, cap-based design (fixed offset, variable inter-sensor distance), rigid generic helmets (variable offset, fixed inter-sensor distance), and an idealised fixed offset, equidistant inter-sensor array.

In all cases the sensor count was fixed at 64 (192 channels; tri-axial) to match our physical system.

#### 2.3.1. Head Geometry

Head geometry was derived from structural MRI images acquired from 24 participants (18 male, 6 female) with a mean age of 40.3 (SD = 12.16) for whom scannercasts had been produced in a previous study, described in related simulation work (Alexander et al., 2025). Note that following exclusion criteria (see *2.3.4. Rigid Helmet Dataset*) the 18 remaining participants (13 male, 5 female) had a mean age of 38.7 (SD = 12.09). The original study was approved by the University College London Research Ethics Committee (project ID 6743/007). We took this approach, rather than using template anatomy, to improve the generalisability of our simulation results across true anatomical variability in head shape and size.

Structural images were defaced and then segmented using unified segmentation (Ashburner & Friston, 2005) as implemented in SPM. The inverse of the ‘other’ tissue class (tissue 6) was then converted into a 3D mesh using the open source iso2mesh toolbox (Fang & Boas, 2009; Tran et al., 2020). The resulting mesh of the scalp was exported as an STL file. Any geometry corresponding to the neck, shoulders or ears was removed manually in Blender.

#### 2.3.2. Sensor Module Dataset

Sensor module arrays were generated for each head model using the CAD software Blender. Individual scalp meshes were imported and a conformal cap with uniform thickness of 3 mm (approximating the neoprene cap) was created for each model (**Figure 1C**). Digital models of the sensor modules, incorporating the location and orientation of individual sensors, were then placed manually on the cap surface.

Module placement was guided by Blender’s snapping tools to ensure realistic positioning, with the aim of maximising spatial density while maintaining feasible mounting. Modules were placed under the same 16 module arrangement, regardless of head size, with reference to fiducial positions (inion, nasion, pre-auricular points).

This mirrors how the caps were physically placed on the head in our empirical datasets.

#### 2.3.3. Equidistant dataset

An equidistant sensor dataset was generated to match the scalp coverage of the sensor module configuration, representing an idealised array equivalent to a 64 sensor scannercast. Because the scalp has non-zero Gaussian curvature, perfect equidistant sampling of its shape is not possible. Instead, an approximation was obtained.

First, the coverage of each sensor module array was quantified relative to anatomical landmarks (left and right pre-auricular, nasion, inion). Sensors were projected into a 2D topographical representation using anatomically veridical projection methods (Alexander et al., 2025) and the convex hull of this projection was computed. This represented the normalised extent and shape of the overall array with respect to the individual head model. A hexagonal lattice with intrinsically equidistant vertices was then overlaid onto this hull such that 64 vertices lay within the boundary. Finally, the lattice was scaled to match the surface area of the original coverage. The result of this can be seen in **Figure 2B**.

**Fig. 2.**
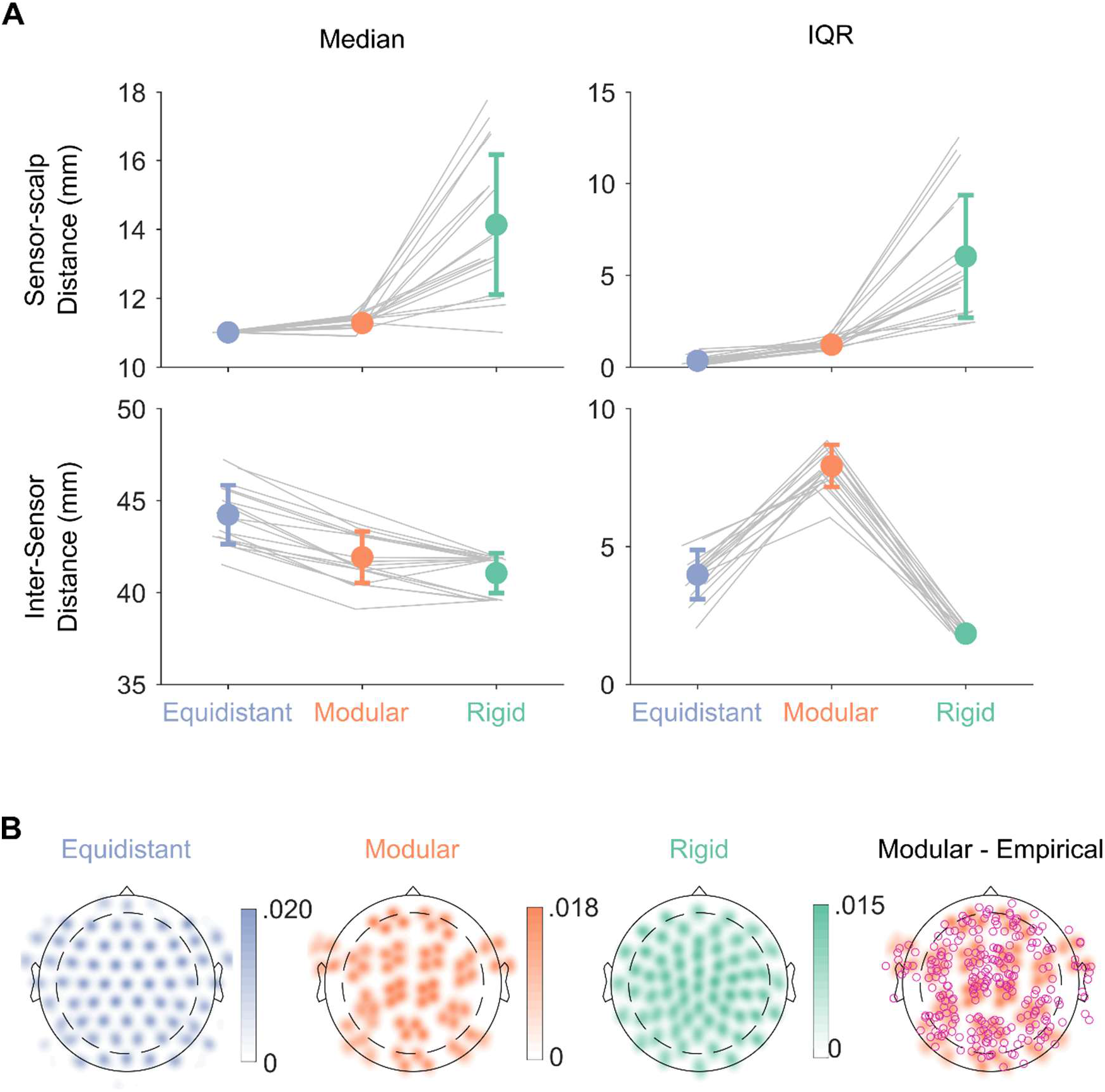
Sensor array properties under the three simulated sensor array conditions. (A) Description of sensor positioning with respect to the scalp and other sensors. Top panels show the distribution of sensor-scalp distances as median and inter-quartile range (IQR) in each array condition (equidistant, modular, rigid). Bottom panels show inter-sensor distances. Grey lines connect data points from the same head shape. Error bars show the standard deviation about the mean. (B) Sensor coverage shown as smoothed 2D density plots of sensors, collapsed across head shapes within array conditions. Smoothing was achieved using a Gaussian kernel (σ = 0.03, image size 1.5 x 1.5). On the right, sensor positions from our five empirical datasets are overlaid as pink circles on top of the simulated map.

These 2D positions were then projected back into 3D using the inverse anatomical projection, with sensors’ positions at a fixed offset of 11 mm from a convex hull of the scalp surface. This served as a starting point to produce an inter-sensor equidistant array. The neighbourhood between positions was defined by the original lattice structure. Median inter-sensor distances were calculated for each sensor across these connections.

Equidistance was further refined iteratively. A density map was generated by interpolating average distances (in 3D) across the 2D topography. The gradient of this map was then used to compute a weighted descent vector for each sensor, yielding updated 2D positions. These were projected back into 3D and re-evaluated. Iteration continued until further updates increased the range of median inter-sensor distances.

Next, tri-axial orientations were created. For each sensor position a radial vector from the head origin (defined as the midpoint between left and right pre-auricular) was first taken. To define the primary tangential axis, a geodesic curve on the scalp surface passing through left pre-auricular, the sensor position and right pre-auricular was made and its local Euclidean direction at the sensor position was taken as an initial tangential estimate. This initial vector was then orthogonalized with respect to the radial direction. The secondary tangential axis was then defined as the cross product of the radial and primary vectors, completing an orthonormal tri-axial frame for each sensor.

#### 2.3.4. Rigid Helmet Dataset

A commercially available mobile mounting system, compatible with the Neuro-1, is produced by Cerca Magnetics (Cerca Magnetics Ltd., Nottingham, UK). These are generic rigid helmets are available in different sizes, each with 64 sensor slots. For our simulations we used the small and large adult sizes only. The helmets comprise a single rigid structure such that the whole sensor array is positioned as one. We selected these helmets for comparison as they have been demonstrated to be effective at measuring MEG signals (Brookes et al., 2022; Hill et al., 2022).

To digitally replicate realistic helmet placement, we applied a rigid body transform to 3D models of the small and large adult helmet sizes, aiming to achieve a close fit to the head while preventing geometric overlap between the head and helmet meshes. This was implemented using a custom Matlab tool. Placement was performed manually using a graphical user interface in which the user (N.A.A.) adjusted translation and Euler rotation via sliders. By doing so, the helmet was positioned such that the eyes remained unobstructed and the internal helmet surface contacted the scalp at the vertex.

This procedure was repeated for both helmet sizes. When the small helmet fit, it was used (6/24). Otherwise, the large helmet was used (12/24). When neither size fit (6/24) the corresponding head geometry was excluded from all simulated conditions since no commercially available helmet could be selected.

The rigid body transform applied to the helmet was then applied to the corresponding sensor positions and orientations.

#### 2.3.5. Leadfield Simulations

For the 18 head geometries which had a complete set of three array configurations we produced individual cortical meshes using non-linear transformation of the MNI canonical 20,248 vertex cortical mesh in SPM. For each geometry-array pairing we then calculated leadfields using the Nolte single shell forward model (Nolte, 2003) implemented in Fieldtrip (Oostenveld et al., 2011).

#### 2.3.6. Leadfield Attenuation

To estimate signal loss under each of the array configurations, we calculated leadfield attenuation for the modular and rigid arrays relative to the idealised fixed offset, equidistant sensor array. Specifically, the leadfield attenuation for each participant (*s*) at each vertex index (*i*) was defined as

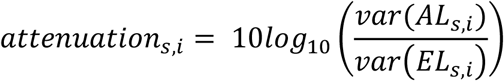

Where *AL* was the leadfield for the modular or rigid array and *EL* was the leadfield for the equidistant array. The mean and standard deviation of the attenuation was then calculated across participants, providing one value per array type (modular, rigid), per vertex.

This metric matches that used in Tierney et al. (2020, 2022) to assess signal loss caused by interference suppression techniques. Here we have adapted it to assess the effect of different array designs and also to consider variation between head geometries. By comparing modular and rigid leadfields to the equidistant array leadfield within the same participant’s anatomy, we have controlled for cross-participant anatomical variability while preserving between-array differences in leadfield power.

### 2.4. Empirical Data

To demonstrate the functionality and feasibility of our modular, cap-based OP-MEG system, we collected empirical data from participants performing the same task, either with or without movement.

#### 2.4.1. Participants

Five participants (3 male, 2 female), with a mean age of 32.6 (SD = 6.41), participated in the study. All participants provided written informed consent before the experiment. All passed eligibility criteria for 3T MRI and MEG. The study was approved by the University College London Research Ethics Committee.

#### 2.4.2. General Procedure

Data were collected within a magnetically shielded room (MSR) with internal dimensions 4.4 m x 3.4 m x 2.2 m. To minimise remnant magnetic fields and ensure optimal operating conditions for the OPMs, the MSR was degaussed before each session (Altarev et al., 2015).

MEG data were acquired using a QuSpin Neuro-1 system. Due to ongoing replacement, one sensor out of 64 was unavailable to place on the head. The remaining 63 sensors (189 channels) were placed into 16 modules in the arrangement shown in **Figure 1E**. Sensors were operated in closed-loop mode and sampled at 1500 Hz.

#### 2.4.3. Array Calibration Procedure

Sensor geometry and gain calibration were performed using an array of independently driven external coils (Borna et al., 2017; Hill et al., 2025). We used the QuSpin HALO system mounted ∼10 cm above the head to do this. Details on this system are available in (Hill et al., 2025).

We performed the array calibration procedure three times. Once at the start of a recording, once after the stationary task was completed, and a final time after the ambulatory task. This involved attaching the HALO device to the cap using three mounting points (**Figure 1F**) and then recording OPM data while the inbuilt coils operated. Trigger information corresponding to coil activation was also saved. The HALO device and mount were worn for the duration of the experiment, remaining fixed in relation to the head.

OPM and HALO operation data were processed in SPM to provide sensor-wise calibration of position, orientation and gain. Additionally, coil operation inputs were bootstrapped to provide a robustness estimation (standard deviation across 200 bootstraps) for each of the estimated calibration parameters. If position robustness was greater than 5 mm, orientation robustness greater than 10 degrees, or gain greater than 0.2, the calibration was marked as bad.

#### 2.4.5. Stationary Task Protocol

Participants were seated at approximately the centre of the MSR. After a quality assurance recording, they completed a binaural roving oddball task.

The auditory roving oddball task was adapted from Garrido et al. (2008), and used for consistency with earlier work where this task has been used as an MEG benchmark (Borna et al., 2017; Cook et al., 2024; Seymour et al., 2021). Auditory tones were presented to both ears simultaneously, at a comfortable level. Each tone lasted 70 ms (5 ms rise and fall time) with an inter-stimulus interval of 430 ms. Initial tone frequency was randomly selected from 50 Hz bins between 500 and 800 Hz. The same frequency tone was presented 1-11 times before switching to a new frequency. Sequence lengths were weighted such that short sequences of 1-2 tones were rare (2.5% each), intermediate sequences of 3-4 tones were more frequent (3.75% each), and longer sequences of 5-11 tones were most common (12.5% each).

Stimuli presentation was controlled by PsychoPy (Peirce, 2009) and presented via air tube earphones which were placed in both ears before the task, with the transducers housed outside the MSR (Etymotic; Lucid Hearing Holding Company, LLC, TX, USA).

#### 2.4.6. Ambulatory Task Protocol

We repeated the roving oddball task under ambulatory conditions, replicating work by Seymour et al. (2021). Participants began the task standing in the centre of the MSR. After the first presentation of tones, they were instructed to move continuously around within the room, including turns. An experimenter was present in the room to move the power and data cables for the Neuro-1 OPM system, as required. Participants were asked to support the air tubes for the earphones to reduce noise from movement and to prevent them falling out.

### 2.5. Sensor Array Metrics

We computed metrics from our empirical and simulated datasets to describe the sensor arrays used in our modular, cap-based system, the rigid generic helmet, and the idealised on-scalp, equidistant array.

#### 2.5.1. Proximity of Sensors to Scalp

We measured the distance from sensor to scalp under each of our three simulated array conditions. This was done by ray-casting from sensor position to the head model. An initial vector of sensor position to the nearest vertex was taken and then further optimised to minimise distance.

#### 2.5.2. Inter-sensor Equidistance

Using the neighbourhood information describing connected sensor positions we calculated the median and inter-quartile range (IQR) Euclidean distance from each sensor to its neighbours, under each of our array conditions. Naturally, the rigid helmet arrays would only vary between helmet sizes (small and large).

#### 2.5.3. Head Coverage Metric

For each head geometry/array combination we projected sensor positions into a common 2D space using anatomically veridical projection methods (Alexander et al., 2025). This projection method represents sensor positions in ratio to anatomical landmarks, matching the 10-10 electrode placement system (Jasper, 1958). Bringing participant’s sensor positions into this common space allowed us to compare them directly. To do so, we first interpolated the positions onto a 600 x 600 binned grid. We then collated these grids within each array configuration and calculated the density (count) at each bin, forming a 2D histogram image. Finally, we smoothed this grid using a Gaussian kernel (*σ* = 0.03, image size 1.5 x 1.5).

### 2.6. MEG Analysis

#### 2.6.1. Spatial Co-Registration

Structural MRI scans were acquired for four of the five participants. For the remaining participant, a close matching structural image was used in its place (Holliday et al., 2003). Scalp surfaces were extracted to match our simulation pipeline (see *2.3.1. Head Geometry*).

Per participant sensor positions were initialised using the known rotation from HALO placement on the head (unregistered sensors are in HALO space), and the centroid of the scalp mesh and sensor array subtracted. Translation and rotation of the sensor array was optimised alongside an ‘inflation’ parameter which projected scalp vertices along their normal, to minimise a distance error term between sensors and the scalp surface.

#### 2.6.2. Pre-processing

The Neuro-1 acquisition software applied a 500 Hz low-pass filter before saving the data. We characterised the delay introduced by this filter as 5 ms. Reported timings have been corrected by applying a corresponding counter-offset.

Calibrated gain was first applied and then bad channels were discarded having been identified using two criteria. First, channels were excluded if reliable sensor calibration could not be estimated (see *2.4.3. Array Calibration Procedure*). Second, channels whose power spectra deviated substantially from the median were identified through visual inspection and excluded. This resulted in a mean of 58.6 (SD = 2.2) good sensors and 167.4 good channels (SD = 7.3) across recordings.

A series of pre-processing steps were applied to the remaining channels. Zapline Plus (de Cheveigné, 2020; Klug & Kloosterman, 2022) was first applied to suppress line noise at 50 and 120 Hz. For one participant, Zapline Plus was also applied at 23.5 Hz to remove interference believed to be caused by signal mixing between the OPM amplitude modulation frequency (Cohen-Tannoudji et al., 1970) and line noise (50 Hz). A 5^th^ order two-pass Butterworth notch filter (574–580 Hz) was then applied to suppress aliased modulation frequency activity, followed by a 5^th^ order two-pass Butterworth band-pass filter (2-80 Hz). An adaptive multipole model (AMM; internal harmonic order = 9; external harmonic order = 2) based filter (Tierney et al., 2024) was then applied with temporal extension (correlation limit = 0.95). Finally, the data were downsampled to 500 Hz.

Continuous data were then segmented into epochs. For roving oddball task trials, a time window was selected from 100 ms before tone onset to 400 ms after tone onset. The random nature of oddball timing resulted in a small variation in trial number with a mean of 594.6 trials (SD = 21.04) in the seated version and a mean of 584.2 trials (SD = 43.48) in the ambulatory equivalent.

For each participant and dataset, bad trials were labelled by visual inspection in Fieldtrip using the method *ft_reject_visual*, to assess the maximum absolute range within each trial. Across participants, an average of 12.8 bad trials (SD = 13.7) were removed from the seated auditory task and 66.6 bad trials (SD = 43.4) from the ambulatory auditory task.

#### 2.6.3. Source Reconstruction

Individual structural MRI images (with the exception of one participant, see *2.6.1. Spatial Co-Registration*) were used to non-linearly warp the canonical 5,124 vertex cortical mesh in SPM to each participant’s anatomy. Source reconstruction was then performed on this common cortical mesh using group inversion (Litvak & Friston, 2008) with the Empirical Bayesian Beamformer (Belardinelli et al., 2012). The number of spatial modes was 76 and the number of temporal modes per participant was 16, based on automatic optimisation. A Hanning window was first applied and the inversion was restricted to good trials only. Data from the auditory roving oddball task were first band-pass filtered from 2-40 Hz and the whole trial window included. Leadfields were pre-multiplied by the AMM projector weights before inversion.

We then reconstructed source activity for each trial individually, using the source model estimated for that participant during group inversion and projecting it through the spatial and temporal modes to yield paired baseline (-40 to 0 ms) and M100 activity images (80 to 120 ms). All images were then smoothed directly on the cortical mesh (*spm_mesh_smooth*, 16 iterations).

#### 2.6.4. Source-level Statistics

We aimed to test whether the auditory evoked field (M100) was detectable in the five datasets collected under both seated and ambulatory movement conditions. A fixed-effects analysis at the trial level allows us to test this within our specific sample, without need to support population-level inference (Friston et al., 1999).

The seated and ambulatory versions of the task were analysed independently. For each, a flexible factorial design was specified in SPM with participant identity as an independent factor and time (activity vs. baseline) as a dependent factor. The effect of time was then estimated and an F-contrast used to test the effect of condition, with vertex-and cluster-level family wise error (FWE) correction.

## 3. RESULTS

### 3.1. Array Properties

Array properties including distance to scalp, inter-sensor distance, and head coverage are fundamental to MEG signal amplitude. We characterised these measures in simulation for 18 head shapes across three array conditions. A summary of simulated array metrics can be seen in **Figure 2**.

#### 3.1.1. Sensor-scalp Offset

To quantify sensor-scalp offset, we computed, for each head shape, the median and interquartile range (IQR) of sensor-scalp distance across the 64 sensors. These summary metrics were then averaged across head shapes to characterise the typical distance and consistency within arrays for each array condition. These results are shown in **Figure 2A**.

The idealised equidistant array showed highly consistent sensor-scalp distances across all head shapes, with mean median distance of 11.0 mm (SD = 0.016 mm) and mean IQR of 0.36 mm (SD = 0.258 mm). Our proposed module arrays were similarly close with mean median distance of 11.3 mm (SD = 0.185 mm), but exhibited a larger range with mean IQR of 1.23 mm (SD = 0.213 mm). The rigid arrays exhibited both larger and substantially more variable sensor-scalp distances with mean median distance of 14.1 mm (SD = 2.032 mm) and mean IQR of 6.03 mm (SD = 3.333 mm)

These results serve as a manipulation check for the equidistant and modular arrays, both of which were designed to maintain 11 mm offset from the scalp. For the rigid array condition, the observed mean distance is consistent with reported values for real application of the generic rigid helmet system (e.g. ∼14.5mm (Rier et al., 2024); ∼21 mm (Rhodes et al., 2023) suggesting our simulated placement procedure was in line with physical implementation.

#### 3.1.2. Inter-sensor Distance

To quantify inter-sensor spacing, we repeated these measures (median and IQR within-array, across head shapes). As expected, the idealised equidistant condition showed regular inter-sensor spacing across head shapes, with a mean median distance of 44.2 mm (SD = 1.60 mm) and a mean IQR of 3.99 mm (SD = 0.89 mm). Our proposed module arrays exhibited slightly reduced spacing and greater variability, with a mean median distance of 41.9 mm (SD = 1.40 mm) and a mean IQR of 7.93 mm (SD = 0.76 mm). In contrast, the rigid arrays showed the smallest inter-sensor distance and the lowest variability, with a mean median distance of 41.1 mm (SD = 1.08 mm) and a mean IQR of 1.85 mm (SD = 0.15 mm). Note that for rigid arrays, only two sets of values were possible, determined by the helmet size being either small or large.

#### 3.1.3. Head Coverage

The head coverage of each array is shown in **Figure 2B**. The idealised and modular arrays have roughly equivalent coverage, when projected into a common 2D space. Module placement was consistent, regardless of large variations in head size, as shown by identifiable clusters of sensors within modules. Variation increased at the periphery. Rigid helmets were also consistent and in line with their intended placement as it aligns with the 10-10 EEG convention (Seedat et al., 2024). This is shown by full coverage within the dashed circumference, which represents the standard extents of this placement system (Jasper, 1958).

#### 3.1.4. Practical Evaluation

The weight of hardware mounted onto the head in addition to the weight of the sensors is a major consideration for any design, particularly where movement and long scanning times are required. A Neuro-1 compatible OPM weighs ∼4 g (256 g for 64 sensors). Each of our modules weighs 14.5 g (232 g for 16 modules). A neoprene cap weighs a further 90 g. Therefore, the total expected weight of our design is 578 g. This compares with 606 g (350 g + sensor weight) for a scannercast and 856 g for a generic rigid helmet (600 g + sensor weight). Therefore, the weight of our module solution is comparable to scannercasts and lighter than an existing commercial solution.

Another important consideration is manufacturing cost. Our modules cost £10 each to print using additive manufacturing methods and neoprene caps were purchased for £250 each. A comprehensive adult system with 22 modules and 8 caps, allowing for fine grained matching to a broad range of individuals, would therefore cost less than a single scannercast (∼£2,500).

Hygiene is a further consideration, particularly for translation into a clinical setting. With our design, only the neoprene caps contact the participant and these can be washed between uses. For paediatric MEG this is of particular importance due to concerns about transmission of head lice.

Finally, we timed the setup of the modules, cap and participant. Each module took approximately one minute to prepare (15-20 minutes total). Removing modules from one cap and transferring them to another size took ∼15 minutes and setting up a participant with the cap and backpack took ∼5 minutes. Naturally, these timings will vary and may change with modifications to the design.

### 3.2. Simulation Results

#### 3.2.1. Signal Amplitude of Brain Sources

To assess how signal amplitude of brain sources was dependent on array type, we first considered leadfield attenuation in our simulated dataset based on 18 head geometries (**Figure 3A**).

**Fig. 3.**
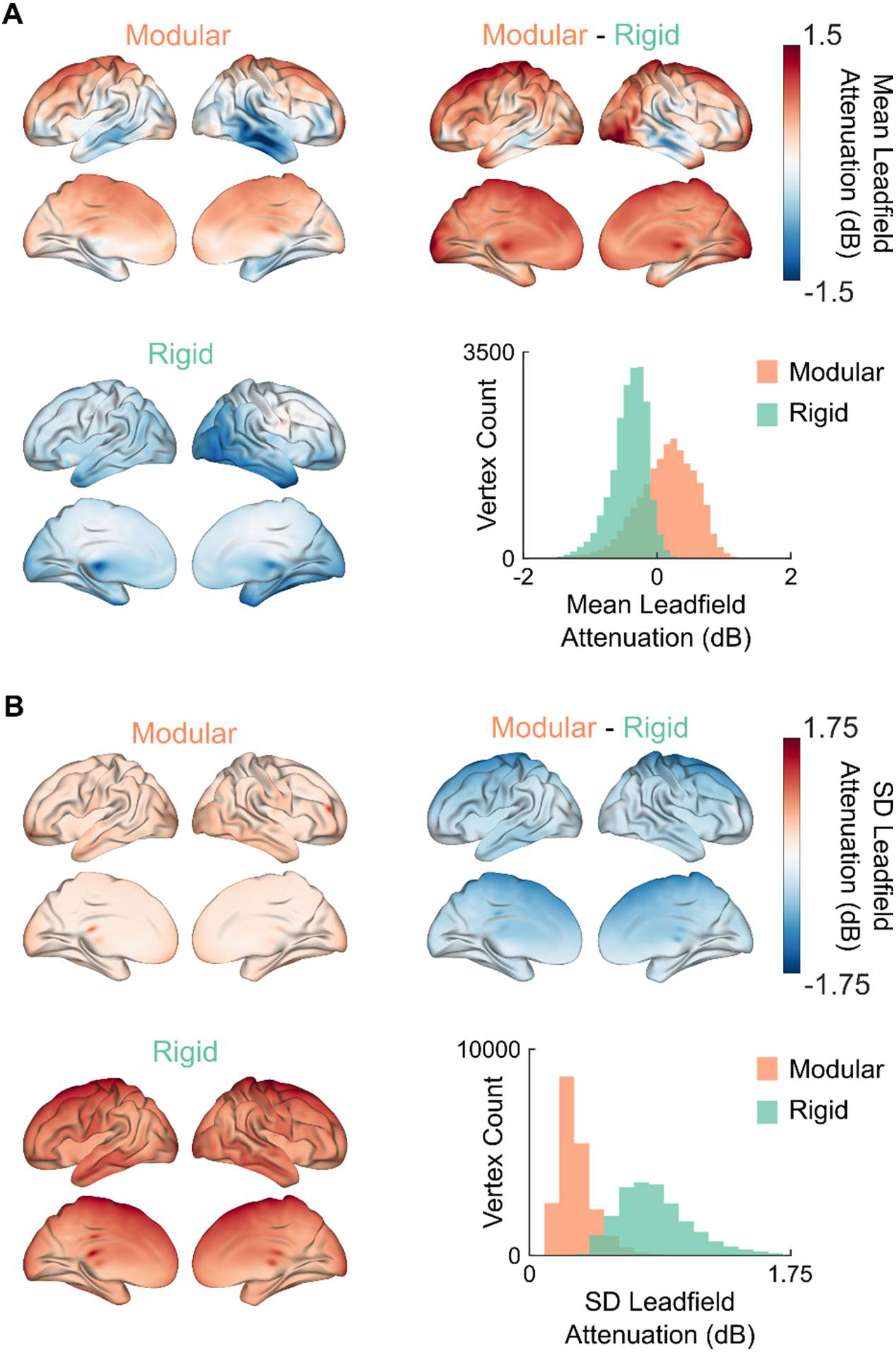
Simulation results showing leadfield attenuation relative to the equidistant array. (A) Mean attenuation (dB), across participants, for modular, and rigid arrays, and their difference. Attenuation values are shown per vertex on an inflated and smoothed mesh for visualisation. The histogram shows the distribution of mean leadfield attenuation relative to equidistant for modular (orange) and rigid (green) arrays. Lower values indicate greater signal loss. (B) The standard deviation (SD) of leadfield attenuation (dB), across individuals, on inflated, smoothed meshes and as a histogram. Higher values indicate greater variation across individuals.

The mean leadfield attenuation for modular arrays relative to the equidistant array baseline, showed a net decrease in leadfield attenuation (mean = 0.14 dB, SD = 0.122), shown by a one-sample t-test against zero, *t*(17) = 4.850, p <.001, d_z_ = 1.143. The spatial pattern showed increased leadfield attenuation for sources in temporal cortex and preserved or slightly reduced leadfield attenuation for frontal and parietal sources. These findings are consistent with the differences in density and head coverage (e.g. no sensors modules placed over the ears).

The rigid arrays showed a net increase in leadfield attenuation (mean =-0.42 dB, SD = 0.601), shown by a one-sample t-test against zero, *t*(17) =-2.962, p =.009, d_z_ =-0.698, particularly for regions in frontal, temporal and occipital regions. This is in line with increased distance to source and reduced density at the periphery of the head and with the array having been placed at the vertex of the scalp.

To compare the modular and rigid approaches we conducted a paired *t*-test of mean leadfield attenuation (i.e. the average across all vertices, within each participant which confirmed a significantly greater attenuation (-0.56 dB) in the rigid condition, *t*(17) = 3.809, p =.001, Cohen’s d_z_ = 0.898.

#### 3.2.2. Variation between Head Geometries

To assess variability across individuals, we compared the standard deviation of leadfield attenuation between modular and rigid arrays (**Figure 3B**). The on-scalp arrangement of sensors meant that modular arrays had limited variation across participants (0.122 dB) and less than that of rigid arrays (0.601 dB), confirmed by a Pitman-Morgan test for equal variances, *t*(16) =-9.479, p <.001.

Taken together, these findings suggest that modular arrays not only preserve more signal on average relative to equidistant coverage, but do so more consistently across individual head geometries than rigid arrays.

### 3.3. Empirical Results

#### 3.3.1. Auditory Evoked Field Measurement

Using a fixed-effects design (Friston et al., 1999) we tested whether auditory evoked fields could be observed using our modular array under seated (stationary) and ambulatory conditions. We examined the M100 response in five participants by first identifying potential clusters at a whole-brain height threshold of p <.001 (uncorrected, F > 10.84) and a minimum cluster-extent of 25 vertices (**Figure 4A**). In what follows the nearest MNI coordinate to the peak vertex of significant FWE-corrected clusters is reported together with the F statistic for that vertex.

**Fig. 4.**
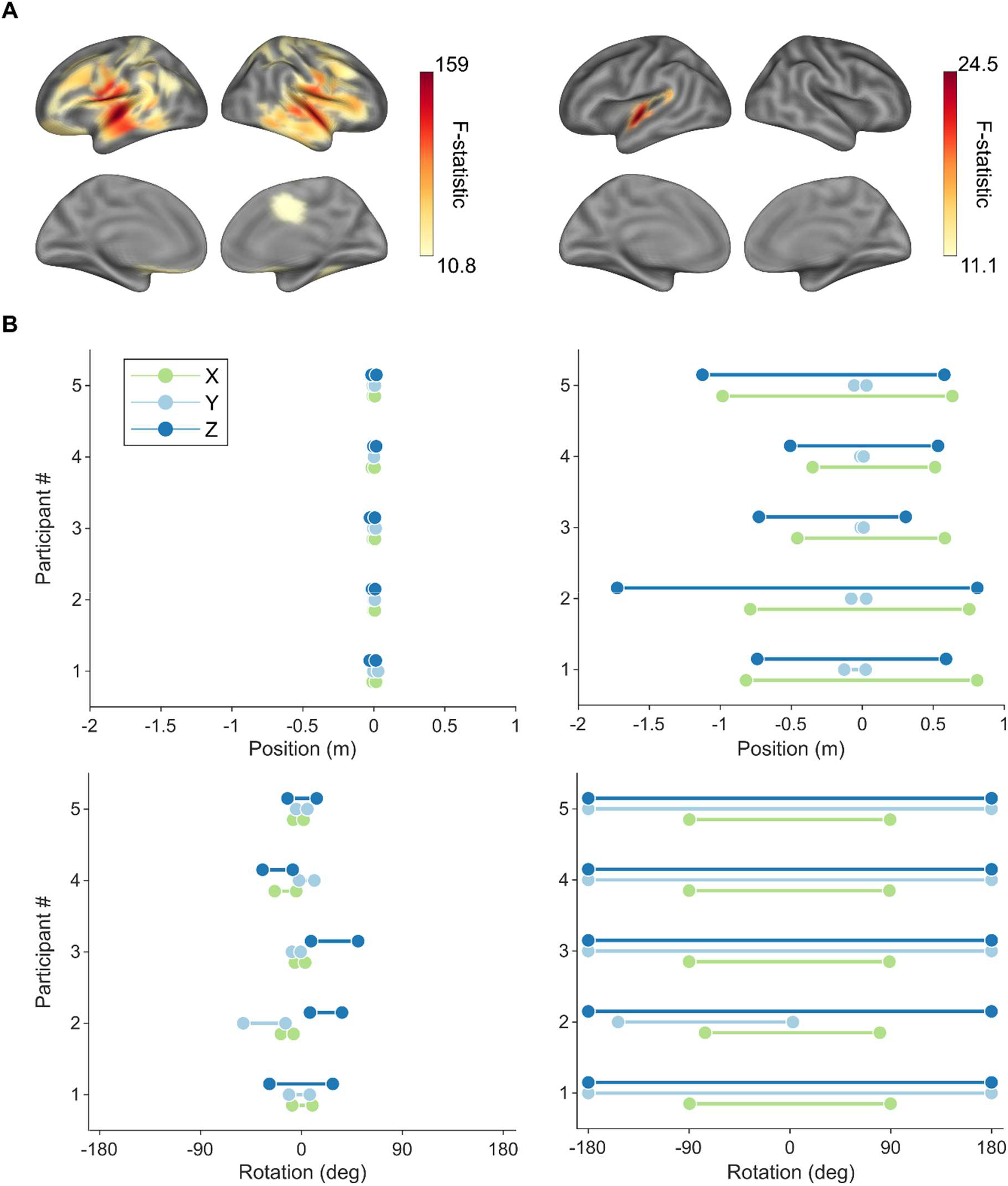
Auditory evoked fields measured under seated (stationary, left column) and ambulatory (right column) conditions. (A) M100 versus baseline activity F-statistic with a mask applied at the vertex-wise, uncorrected p <.001. Data are shown on an inflated mesh and have been smoothed for visualisation. (B) A summary of the range of movement under seated and ambulatory conditions. Translation in X (forward/back), Y (up/down) and Z (left/right) is shown in the upper row and rotation about these axes is shown in the bottom row.

In the seated condition, the response was bilateral with a cluster covering left auditory cortex, MNI [-57,-12,-2], F = 158.80, p < 0.001 (FWE corrected), and right auditory cortex [53,-5,-2], F = 121.18, p < 0.001 (FWE corrected). These clusters were the largest surviving, with the left response slightly larger than the right.

In the ambulatory condition, only the left auditory response survived FWE correction, MNI [-57,-8,-2], F = 24.47, p < 0.001 (FWE corrected). To explore whether a sub-threshold right auditory cortex peak was present, we reduced the cluster forming threshold to p < 0.05. A mirrored pattern of activity was found in the right hemisphere with a cluster in right auditory cortex, MNI [54,-19,-1], F = 6.22, p = 0.013, but it did not pass our main test.

Taken together, these results demonstrate measurement of the M100 event related field, in this sample and under both movement conditions, noting the expected reduction of signal amplitude in the moving condition. Given the robust identification of the M100 response in the stationary condition, this reduction is attributable to movement through background magnetic field gradients, and potentially irregular sound induced by movement of the air tube earphones as a result, rather than the modular array design per se.

#### 3.3.2. Movement Extent

We analysed the six degrees of freedom rigid body motion tracking data collected from the head of each participant during the seated and ambulatory tasks. A visual summary of the movement extent is provided in **Figure 4B**.

When seated, the mean range of movement in the forward and backward axis was 0.016 m (SD = 0.005), in the left and right axis was 0.031 m (SD = 0.010), and in the up/down axis was 0.016 m (SD = 0.011). The average speed of participants was 0.0029 m/s (SD = 0.0005) confirming that participants were effectively stationary.

During the ambulatory auditory task, the mean range of movement in the forward and backward axis was 1.34 m (SD = 0.36 m), in the left and right axis was 1.53 m (SD = 0.63 m), and in the up/down axis was 0.077 m (SD = 0.055 m). The average speed of participants was 0.161 m/s (SD = 0.059 m).

#### 3.3.3. Sensor and Anatomy Spatial Co-registration

As for the simulated data, we calculated the sensor-scalp and inter-sensor distances for each empirical *in vivo* dataset. Across participants, the mean median sensor-scalp distance was 7.40 mm (SD = 1.54 mm) with a mean IQR of 4.72 mm (SD = 1.15 mm).

This was lower than the expected 11 mm. First, compression of the cap and soft tissue on the scalp can reduce the spacing. Second, the method used to produce the scalp models used in simulation and empirical co-registration may apply a small amount of unintended scaling. Third, the assumed sensor to housing offset (6.25 mm) may be inaccurate due to manufacturing tolerances for one or both of the modules and sensors. Importantly, our spatial co-registration accounts for these sources of error by fitting shape, not distance.

Inter-sensor distances were in line with what was expected from simulation. Across individuals the mean distance was 44.90 mm (SD = 2.94 mm) and the mean IQR was 11.80 mm (SD = 0.18 mm). Note that the number of functioning, calibrated sensor per participant varied and none had the full complement of 64 (mean = 51.2, SD = 3.701). Despite gaps due to sensor failures, head coverage closely matched the simulated arrays (**Figure 2B**).

## 4. DISCUSSION

Optimisation of sensor mounting for OP-MEG is crucial to maximising SNR for wearable systems. Therefore, we present a modular, cap-based design for on-scalp sensor mounting that maintains the unique potential of OPMs: mobility, signal amplitude via proximity to source, affordability, and in a manner conducive to high throughput scanning. The result is a system that can adapt to multiple imaging applications, without compromising signal quality.

Our modular configuration builds on earlier simulation and empirical work that has demonstrated the importance of array characteristics (e.g. proximity to source, density, coverage, stability) alongside sensor specification (e.g. calibration error, noise floor). Proximity to scalp is essential to recover the advantage lost by noisier sensors, when compared with SQUID-based MEG. That said, proximity has the greatest benefit for superficial cortical sources with diminishing increase in performance for deeper sources (Brookes et al., 2022). One way to target deeper sources is to improve sensor coverage. For example, by including temporal and face measurements (Feys et al., 2025; Tierney et al., 2021) to increase hippocampal SNR. Our simulations support these conclusions with a new focus on head shape variability.

Our simulations also served to address a potential limitation of our module design: distances between modules vary across the head and across participants and equidistant sampling is not achieved. Assuming equidistant sampling to be ideal (Ahonen et al., 1993), our simulations explored the relative leadfield attenuation cost of modular placement. Overall, our proposed modular arrays showed comparable signal amplitude across cortex when compared to the idealised, fixed-offset array, and an improvement versus a widely adopted generic helmet configuration. However, signal amplitude in temporal regions was reduced under modular configuration. This can be attributed to absent sensor placement over the ears which is necessary for our cap-based solution, but not for the ideal placement simulated, or the off-scalp helmet tested. Future cap-based designs could incorporate additional coverage, with consideration for delivery of auditory stimuli. Importantly, our modular, cap-based design showed consistently minimal leadfield attenuation across participants versus ideal, equidistant placement, reducing the impact of individual head shape on observations.

A great advantage of on-scalp imaging that is facilitated by our modular, cap-based design is that knowing the positions of the sensors inherently provides knowledge about the shape of the head, simplifying the challenging problem of co-registering sensor space to individual anatomy. While this information is inherently known for scannercasts, it is not known for rigid generic solutions that do not conform to the head. A plethora of methods are available to researchers for co-registration, typically involving spatial imaging of the head with and without the sensors, along with building a model of sensor array geometry, and finally aligning the two (Cao et al., 2023; Rhodes et al., 2023; Zetter et al., 2019). However, these require additional scan time and complexity, which are circumvented by our design.

In this work, we used external coils mounted to the top of the cap (QuSpin’s HALO system) to calibrate out sensors, but future work could employ differently shaped coil arrays more suited to flexible sensor arrays (Xu et al., 2025). External matrix coils have also been shown to be effective for array calibration (Hill et al., 2025). Any solution must be able to accommodate at least small movements to promote participant inclusivity. Another benefit of on-scalp measurement is reduced impact of spatial and orientation jitter on SNR (Iivanainen, 2026) and the assurance of equivalent sensitivity to off-scalp measurements even when jitter is applied (Zetter et al., 2018).

### 4.1. Future Outlook

Sensor count in OP-MEG systems will inevitably increase beyond the 64 sensor (192 channel) system assessed here. This will exacerbate challenges of mounting individual sensors, unless a modular approach is taken. As sensors become smaller our solution can easily be adapted to accommodate more sensors per module. Ideally for the end user, modules following the design principles outlined here would be adapted into the OPM sensor manufacture process, with shared OPM cabling per module.

Beyond MEG, wearable arrays of OPMs, which can be arranged flexibly, create opportunities for biomagnetic measurement extending to the wider central nervous system, the heart, muscles and gut. Crucially, our modular design can achieve this in a unified system, facilitating basic neurophysiological and physiological investigation through interaction (Spedden et al., 2026; Woelk et al., 2026).

As OP-MEG matures its intended role should be continually reviewed. One trajectory aims to replace cryogenically cooled SQUID-based MEG with static, radially adjustable OPM arrays (Alem et al., 2023; Gutteling et al., 2023). The performance of such arrays already equates to or outperforms contemporary SQUID-based MEG systems (Gaetz et al., 2026). However, it also inherits some of the issues of static SQUID-based MEG, that limit clinical adoption (Gaetz et al., 2015), including requiring participants to remain still. A second trajectory for OP-MEG prioritises wearability and mobility. The capacity for individuals to move, intentionally or unintentionally, during scanning is critical for clinical translation and must be permitted without compromising signal quality. Permitting movement and engagement with the environment also expands the breadth of truly naturalistic behaviour that can be studied with OP-MEG. More ambitious still is the goal of operating OP-MEG beyond the confines of a magnetically-shielded room (Bezsudnova et al., 2024; Limes et al., 2020; Zhang et al., 2020; Zheng et al., 2026), to maximise research opportunities and the clinical translation possibilities to enhance patient care. The modular, cap-based design we have presented here can add benefit to each of these future scenarios by providing a comfortable, low-cost, hygienic method of reproducible on-scalp sensor placement that is robust to movement, reduces analysis complexity, and enables rapid, high throughput imaging and large cohort studies.

## Author Contributions

Conceptualisation: N.A.A., A. M., S.P., Y.B., T.M.T., G.R.B., M.F.C. Software; N.A.A., A. M., S.P., Y.B., T.M.T., G.R.B. Writing - Original Draft: N.A.A., M.F.C. Writing - Review & Editing: A. M., S.P., Y.B., T.M.T., G.R.B., M.F.C. Visualisation: N.A.A., M.F.C. Supervision: T.M.T., G.R.B., M.F.C. Funding acquisition: N.A.A., G.R.B., M.F.C.

## Acknowledgements

We would like to thank Sienna Griffin-Shaw and Felix Roberts at the Bartlett Faculty of the Built Environment, UCL, for their advice regarding additive manufacturing. We would also like to thank Joanne Alexander for advice on cap fabrication.

